# Adventive larval parasitoids reconstruct their close association with spotted-wing drosophila in the invaded North American range

**DOI:** 10.1101/2021.12.06.471517

**Authors:** Paul K. Abram, Michelle T. Franklin, Tracy Hueppelsheuser, Juli Carrillo, Emily Grove, Paula Eraso, Susanna Acheampong, Laura Keery, Pierre Girod, Matt Tsuruda, Martina Clausen, Matthew L. Buffington, Chandra E. Moffat

**Affiliations:** Agriculture and Agri-Food Canada, Agassiz Research and Development Centre, Agassiz, BC, Canada; British Columbia Ministry of Agriculture, Food and Fisheries, Abbotsford, BC, Canada; University of British Columbia, Faculty of Land and Food Systems, Centre for Sustainable Food Systems, Biodiversity Research Centre, Unceded xʷməθkʷəy̓əm Musqueam Territory; British Columbia Ministry of Agriculture, Food and Fisheries, Kelowna, BC, Canada; Systematic Entomology Laboratory, USDA, Smithsonian Institution, Washington, D.C., USA; Agriculture and Agri-Food Canada, Summerland Research and Development Centre, Summerland, BC, Canada

**Keywords:** Unintentional biological control, adventive establishment, *Leptopilina japonica*, *Ganaspis brasiliensis*, *Asobara*, *Drosophila suzukii*

## Abstract

Two species of larval parasitoids of the globally invasive fruit pest, *Drosophila suzukii* (Diptera: Drosophilidae), *Leptopilina japonica* and *Ganaspis brasiliensis* (both Hymenoptera: Figitidae), were detected in British Columbia, Canada in 2019. Both are presumed to have been unintentionally introduced from Asia, however; the extent of their establishment across different habitats with diverse host plants used by *D. suzukii* was unclear. In addition, there was no knowledge of the temporal dynamics of parasitism of *D. suzukii* by these two parasitoids. We repeatedly sampled the fruits of known host plants of *D. suzukii* over the entire 2020 growing season in British Columbia. We documented the presence of *L. japonica* and *G. brasiliensis* and estimated the apparent percentage of *D. suzukii* parasitized. Across a large region of southwestern British Columbia, both *L. japonica* and *G. brasiliensis* were found to be very common across a variety of mostly unmanaged habitats over the entire course of the season (May-October) in the fruits of most host plants known to host *D. suzukii* larvae. The two parasitoids were responsible for more than 98% of *D. suzukii* larval parasitism and usually co-existed. Parasitism of *D. suzukii* was variable among hosts plants and sites (0-66% percent parasitism) and appeared to be time-structured. Our study demonstrates that the close association between the two larval parasitoids and *D. suzukii* that exists in Asia has evidently been reconstructed in North America, resulting in the highest parasitism levels of *D. suzukii* yet recorded outside of its area of origin.

## Introduction

Populations of highly mobile and polyphagous, exotic insect pests are challenging to manage because the temporal and spatial reach of most pest management tactics is limited and does not extend beyond cultivated hosts (Rabb 1978; Kennedy & Storer 2000). To effectively manage such problematic invaders, area-wide pest management strategies which can regulate pest populations across landscapes, host and seasons deliver optimal control are needed. Natural enemies that are able to disperse and attack the pest over a diverse range of habitats and host plants can yield long-term landscape-wide pest population suppression (Schellhorn et al. 2015); however, the degree to which pests are regulated can depend heavily on ecological context. For exotic invasive pests, natural enemies from the pest’s area of origin may be imported and introduced intentionally (‘importation’ or ‘classical’ biological control; Heimpel and Mills 2017) or adventive populations can establish as a result of accidental introductions (‘unintentional’ biological control; Weber et al. 2021). Following the establishment of the introduced natural enemies (regardless of whether they are intentional or adventive), evaluating the extent to which they are attacking the pest across diverse host plants and habitats can inform their potential to provide landscape-wide pest population suppression. Any significant spatial or temporal refuges for the pest that reduce overall levels of natural enemy attack – for example, the pest being able to sustain large populations on particular host plants where it is not attacked by parasitoids – could impact the pest’s abundance and point to a natural enemy having less promise for providing significant pest suppression in different ecological contexts (Hawkins et al. 1993; Mills and Getz 1996).

Spotted-wing drosophila, *Drosophila suzukii* (Matsumura) (Diptera: Drosophilidae) is an exotic invasive agricultural pest that has presented enormous management challenges in areas where it has established, in large part due to its mobility and polyphagy (reviewed in Tait et al. 2021). In addition to laying its eggs in ripening fruit in cultivated cropping systems, *D. suzukii* also reproduces in the fruits of a wide range of non-crop wild host plant species in natural and semi-natural environments (Lee et al. 2015; Kenis et al. 2016; Thistlewood et al. 2019). Non-crop habitats serve as important sources of populations of *D. suzukii* that disperse to and impact cultivated fruit crops (Tonina et al. 2018; Urbaneja-Bernat et al. 2020), meaning that some component of crop protection strategies should ideally extend beyond the cultivated hosts. Because of its potential to act on *D. suzukii* populations at the landscape-level and season-long scales, biological control became a major focus of research early in the development of management tactics for *D. suzukii* (Wang et al. 2020).

Foreign exploration for specialized, candidate importation biological control agents of *D. suzukii* in its area of origin (China, Korea, and Japan) found that there were two species of larval parasitoids that most commonly attacked *D. suzukii*: *Ganaspis brasiliensis* Ihering (Hymenoptera: Figitidae) and *Leptopilina japonica* (Novković & Kimura) (Hymenoptera: Figitidae) (Daane et al. 2016; Girod et al. 2018a; Giorgini et al. 2019). After considerable research to characterize the specificity and basic biology of these two species (e.g., Girod et al. 2018b; Wang et al. 2018; Wang et al. 2019; Biondi et al. 2021; Daane et al. 2021), *G. brasiliensis* became the leading candidate for importation biological control of *D. suzukii* in North America and Europe (reviewed in Wang et al. 2020).

However, recent unintentional introductions of these larval parasitoids have evidently occurred in restricted regions of North America (*L. japonica* and *G. brasiliensis* in British Columbia, Canada; Abram et al. 2020) and Europe (*L. japonica* in Italy; Puppatto et al. 2020). The extent of their establishment across habitats and different host plants in their new adventive regions is unclear (Abram et al. 2020). In addition, the seasonal dynamics of larval parasitism of *D. suzukii* remain mostly unknown, even in their native range, as studies in Asia were conducted during a limited seasonal time period (Wang et al. 2020). Their adventive establishment in new geographic areas provides an opportunity to characterize the extent of their association with *D. suzukii* under new environmental and evolutionary contexts, which could inform their potential for effective biological control of *D. suzukii* via redistribution or introduction to other areas of the world.

In this study, we explore the associations between *D. suzukii*, its natural enemies, and host plants across a range of habitat types in south coastal British Columbia, Canada. Specifically, we aimed to characterize the extent of *G. brasiliensis* and *L. japonica* establishment and obtain preliminary estimates of parasitism levels of *D. suzukii* over an entire growing season, across a variety of habitats and host plants, in the region where they were first detected in North America. This is the first study to characterize the temporal dynamics of larval parasitism of *D. suzukii* by these parasitoids over an entire growing season. We also conducted more limited surveys in the adjacent Okanagan Valley to determine the geographic extent of parasitism by *L. japonica* and *G. brasiliensis* populations on *D. suzukii* in our study region.

## Materials and Methods

### Focal site collections

To determine the temporal dynamics of larval parasitism of *D. suzukii*, six focal collection sites were selected from the lower mainland of British Columbia (Figure 1; Table 1). A range of habitat types were selected, including: forest (low-or mid-elevation), a semi-urban park, a community garden, a wetland, and mixed agriculture on an experimental farm (a mixture of wild host plants and unsprayed experimental berry fields). Host plants across these sites included a mixture of cultivated plants: e.g., strawberries, *Fragaria x ananassa* Duchesne (Rosaceae); raspberries *Rubus idaeus* L. (Rosaceae); blackberries, *Rubus fruticosis* L. (Rosaceae); highbush blueberries, *Vaccinium corymbosum* L. (Ericaceae);native non-cultivated plants:e.g., salmonberry, *Rubus spectabilis* Pursh (Rosaceae); red elderberry, *Sambucus racemosa* L. (Adoxaceae), and invasive berries i.e. Himalayan blackberry, *Rubus armeniacus* Focke (Rosaceae) (see Table 1 for full list). Sites were sampled from late May to late August or mid-September, with the exception of the Abbotsford community garden site that was sampled until the end of October, where *R. armeniacus* fruit remained later in the season than at the other sites.

**Table 1.** Parasitism of *D. suzukii* (*Ds*) by larval parasitoids at six focal sampling sites in the lower mainland of British Columbia, Canada, when fruit was repeatedly sampled from different host plant species.

| Site | Host plant species | Total N<br>fruits | Number of<br>sampling dates<br>(date range) | Total<br>*Ds*<sup>a</sup> | % parasitism<sup>b</sup> |  |  | % parasitoid species<br>composition<sup>c</sup> |  |  |  |
| --- | --- | --- | --- | --- | --- | --- | --- | --- | --- | --- | --- |
|  |  |  |  |  | Total | Min | Max | *Lj* | *Gb* | *As* | *Sp* |
| Delta<br>(semi-urban<br>park) | *Rubus spectabilis* | 146 | 5 (Jun 11-Jul 9) | 211 | 7.0 | 0.0 | 30.4 | 43.8 | 56.3 | 0.0 | 0.0 |
|  | *Sambucus racemosa* | 8,968 | 3 (Jun 25-Jul 9) | 83 | 53.0 | 48.0 | 64.3 | 0.0 | 100.0 | 0.0 | 0.0 |
|  | *Rubus armeniacus* | 155 | 5 (Jul 23-Aug 27) | 134 | 3.6 | 0.0 | 18.2 | 0.0 | 100.0 | 0.0 | 0.0 |
| Maple Ridge<br>(low-elevation<br>forest) | *Rubus spectabilis* | 130 | 5 (Jun 3-Jul 1) | 305 | 7.0 | 0.0 | 24.6 | 100.0 | 0.0 | 0.0 | 0.0 |
|  | *Sambucus racemosa* | 1,720 | 3 (Jun 17-Jul 9) | 92 | 8.0 | 0.0 | 22.2 | 12.5 | 87.5 | 0.0 | 0.0 |
|  | *Vaccinium parvifolium* | 420 | 5 (Jun 17-Jul 16) | 201 | 12.6 | 0.0 | 40.3 | 24.1 | 75.9 | 0.0 | 0.0 |
|  | *Oemleria cerasiformis* | 80 | 2 (Jun 19-Jul 1) | 53 | 1.9 | 0.0 | 5.0 | 100.0 | 0.0 | 0.0 | 0.0 |
|  | *Rubus idaeus* | 150 | 5 (Jul 9-Aug 6) | 271 | 24.3 | 11.1 | 64.3 | 47.1 | 50.6 | 2.3 | 0.0 |
|  | *Gaultheria shallon* | 135 | 3 (Jul 30-Aug 14) | 80 | 0.0 | 0.0 | 0.0 | --- | --- | --- | --- |
|  | *Rubus armeniacus* | 120 | 4 (Jul 31-Aug 27) | 451 | 12.9 | 0.0 | 29.9 | 92.5 | 7.5 | 0.0 | 0.0 |
|  | *Prosartes hookeri* | 75 | 2 (Aug 14-Aug 21) | 69 | 4.2 | 0.0 | 11.1 | 100.0 | 0.0 | 0.0 | 0.0 |
| Abbotsford<br>(community<br>garden) | *Sambucus racemosa* | 41,822 | 9 (Jun 5-Jul 31) | 1071 | 22.2 | 6.6 | 55.4 | 40.2 | 59.8 | 0.0 | 0.0 |
|  | *Rubus armeniacus* | 428 | 13 (Jul 24-Oct 27) | 1472 | 11.5 | 0.0 | 34.7 | 57.1 | 41.9 | 0.0 | 1.0 |
| Chilliwack<br>(wetland) | *Rubus spectabilis* | 120 | 4 (Jun 2-Jul 2) | 168 | 14.7 | 0.0 | 31.8 | 89.7 | 10.3 | 0.0 | 0.0 |
|  | *Sambucus racemosa* | 2,020 | 3 (Jun 15-Jul 2) | 245 | 21.5 | 5.3 | 56.0 | 34.3 | 65.7 | 0.0 | 0.0 |
|  | *Rubus parviflorus* | 90 | 2 (Jul 2-Jul 10) | 41 | 4.7 | 0.0 | 9.1 | 100.0 | 0.0 | 0.0 | 0.0 |
|  | *Rubus armeniacus* | 210 | 7 (Jul 24-Sep 7) | 1092 | 15.5 | 0.0 | 35.2 | 71.1 | 27.9 | 1.0 | 0.0 |
| Chilliwack<br>(mid-elevation<br>forest) | *Rubus spectabilis* | 150 | 5 (May 28-Jul 2) | 53 | 1.9 | 0.0 | 6.7 | 100.0 | 0.0 | 0.0 | 0.0 |
|  | *Sambucus racemosa* | 3,820 | 6 (Jun 22-Jul 30) | 264 | 34.8 | 0.0 | 65.6 | 7.8 | 92.2 | 0.0 | 0.0 |
|  | *Oemleria cerasiformis* | 180 | 3 (Jun 22-Jul 9) | 10 | 0.0 | 0.0 | 0.0 | --- | --- | --- | --- |
|  | *Rubus parviflorus* | 195 | 4 (Jul 16-Aug 7) | 121 | 9.0 | 0.0 | 16.3 | 100.0 | 0.0 | 0.0 | 0.0 |
|  | *Rubus armeniacus* | 150 | 5 (Aug 7-Sep 7) | 1154 | 15.9 | 2.0 | 34.4 | 67.9 | 32.1 | 0.0 | 0.0 |
| Agassiz<br>(experimental<br>farm) | *Fragaria x ananassa* | 168 | 7 (Jun 2-Sep 16) | 308 | 27.2 | 0.0 | 45.0 | 93.0 | 7.0 | 0.0 | 0.0 |
|  | *Sambucus racemosa* | 2,631 | 5 (Jun 7-Jul 8) | 92 | 3.2 | 0.0 | 4.1 | 0.0 | 100.0 | 0.0 | 0.0 |
|  | *Rubus idaeus* | 150 | 5 (Jun 24-Jul 22) | 710 | 27.9 | 0.0 | 60.0 | 83.3 | 16.4 | 0.4 | 0.0 |
|  | *Vaccinium corymbosum* | 630 | 7 (Jul 1-Aug 12) | 658 | 1.8 | 0.0 | 5.7 | 100.0 | 0.0 | 0.0 | 0.0 |
|  | *Rubus armeniacus* | 180 | 6 (Jul 25-Aug 26) | 1204 | 12.1 | 0.0 | 39.8 | 95.8 | 4.2 | 0.0 | 0.0 |
|  | *Rubus fruticosus* | 240 | 8 (Jul 25-Sep 16) | 3310 | 18.2 | 0.0 | 54.3 | 82.4 | 16.1 | 1.5 | 0.0 |

<sup>a</sup> Estimate of total number of *Drosophila suzukii* pooled among samples, including parasitized individuals (=emerged flies + emerged parasitoids).
<sup>b</sup> Total parasitism– pooled among all samples; min/max parasitism – minimum and maximum values across all sampling dates.
<sup>c</sup> Percentage of parasitoids of each species, calculated with data from all sampling dates pooled. *Lj* - *Leptopilina japonica*; *Gb* - *Ganaspis brasiliensis*; *As* - *Asobara* cf. *rufescens*.; *Sp* – *Spalangia erythromera*.

**Figure 1.**
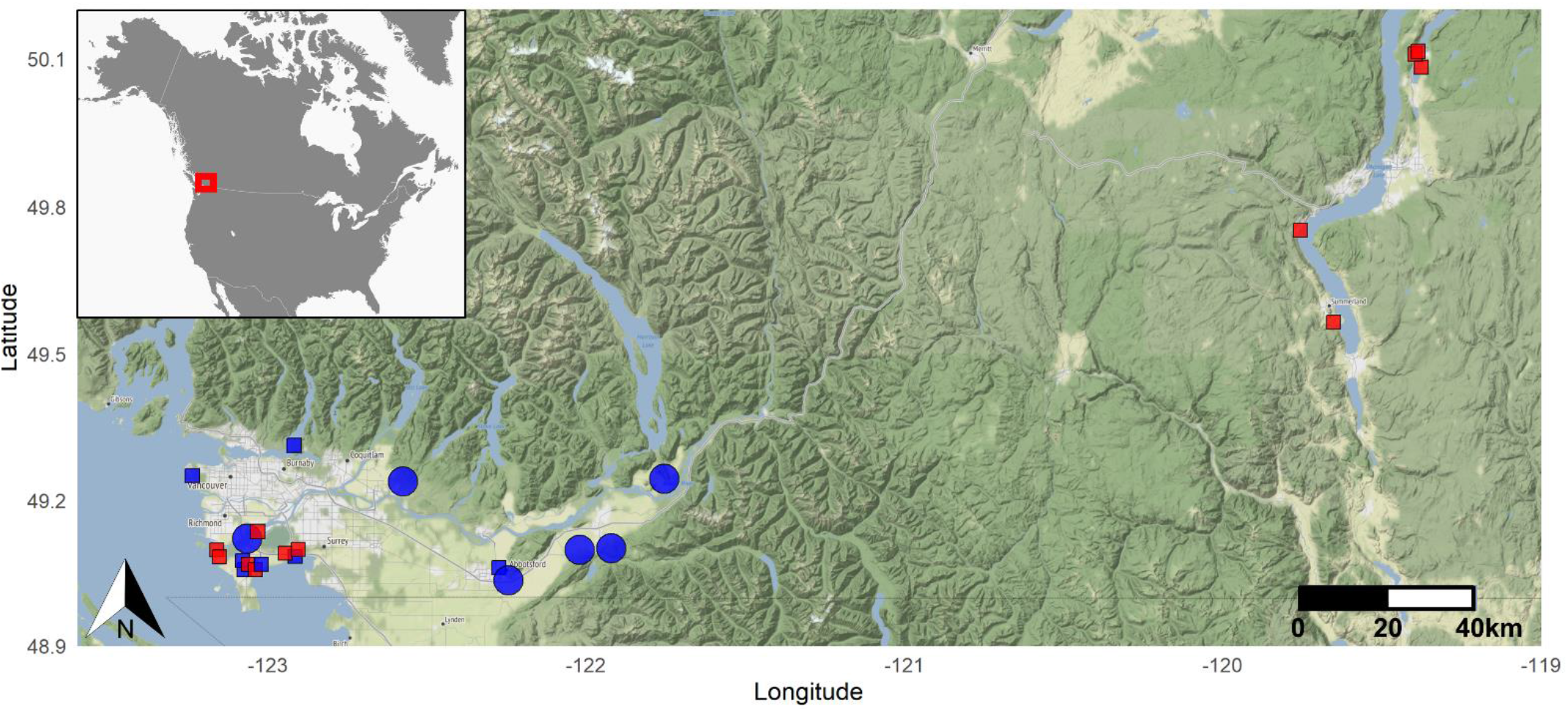
Map of locations where adventive larval parasitoids of *Drosophila suzukii* were and were not detected in British Columbia, Canada in 2020 (only sites where *D. suzukii* were reared from fruit are shown). **Large circles** - focal sampling sites with season-long sampling; **small squares** - non-focal sampling sites with limited sampling. **Blue symbols** - both *Leptopilina japonica* and *Ganaspis brasiliensis* found; **red symbols** - no larval parasitism found. The red box within the inset shows where the study area is located in North America. Map tiles by Stamen Design, under CC BY 3.0. Data by OpenStreetMap, under ODbL.

All focal sites surveyed had (i) a minimum of two host plant species collected from at least two collection dates and (ii) three fruit samples, referred to herein as “pseudoreplicates” collected from each of the two collection dates. Fruit collections began when the first ripe fruit were available to pick and ended in cultivated crops two weeks following the last harvest and in wild fruit when fruit was scarce and remaining fruit was rotten or dried up. Sampling intervals during the fruiting period ranged from every 4-12 days but were typically once per week (see Table S1). Standardized host plant sampling collection plans were designed to encompass the spatial variation in the arrangement of host plants. As our sites were pre-existing, the arrangement of host plants was not consistent. We thus categorized the spatial arrangement of host plants as follows: (1) *rectangular* field divided into rows, (2) *linear* patches accessible from one side, (3) *large polygonal* patches accessible from both sides, and (4) *multiple small polygonal* patches or individual plants (see Abram et al. 2022, Figure 4). For a rectangular patch, fruit sampling locations were chosen by randomly selecting three rows and sampling positions within those rows. For a linear or large polygonal patch, the first sampling point was randomly selected within the first 25% of the total linear or perimeter distance of the patch, with the two subsequent sample locations adding 25% of the total patch length. When multiple small patches or individual plants were sampled, three patches were randomly selected from the available patches and a random starting point was chosen. For all patch types, fruits were collected from within a 10 m section for each selected sampling location and fruit sampling locations were selected anew each week.

Due to the differences in fruit size among host plants, we varied the number of berries collected by the host plant to achieve a similar total fruit weight (typically ∼20-60 g) for each host plant for each collection. For example, we aimed to collect a total of 24 strawberries, 30 raspberries, and 90 blueberries at each site weekly (See Table S1). Fruit collections were held in ventilated rearing containers 12 × 12 × 8 cm (Ziploc Medium Square Container) as described in Abram et al. (2022), but without the wire mesh insert. Fruit samples were monitored three times per week for emergence of *D. suzukii* and parasitoids *L. japonica* and *G. brasiliensis* held at three indoor locations, where mean daily temperatures ranged from 15 to 25°C (Chilliwack site: 15°C, 25°C, 22°C; Maple Ridge site: 15°C, 23°C, 19°C; Abbotsford site: 19°C, 21°C, 20°C; minimum, maximum, and mean temperatures, respectively). In laboratory studies, *Drosophila suzukii* has been found to complete larval development at constant temperatures from 11.8 to 27.5°C and larval parasitoids *L. japonica* and *G. brasiliensis* at temperatures ranging from 17.2 to 27.5°C (Hougardy et al. 2019). Incubation temperatures at our sites were typically within a range suitable for *D. suzukii* and parasitoid larval development and would not be expected to trigger parasitoid diapause (see also below for evidence that parasitoids did not enter diapause). Mean daily temperatures at the Chilliwack and Maple Ridge incubation sites only went below 17°C for 2 and 11 days, respectively.

### Non-focal site collections

To establish the distribution of these two larval parasitoids in areas of BC that were located outside of the focal collection region, we completed single-time point and non-standardized collections of ripe berries from 11 hedgerow sites with non-crop host plants (e.g. *R. armeniacus*; Pacific serviceberry, *Amelanchier alnifolia* [Nuttall] [Rosaceae]) in Delta, cultivated and wild host plants (e.g. *R. idaeus*; *R. armeniacus*; thimbleberry, *Rubus parviflorus* Nuttall [Rosaceae], *V. corymbosum*) from the University of British Columbia (UBC) Farm, a wide range of wild and cultivated host plants from the Summerland Research and Development Centre (Agriculture and Agri-Food Canada) (e.g., sweet cherry, *Prunus avium* L. [Rosaceae]; barberry, *Berberis vulgaris* L. [Berberidaceae]; Oregon grape, *Mahonia aquifolium* [Pursh]), and cultivated and wild host plants (e.g. *P. avium* and *M. aquifolium*) around the city of Kelowna (Figure 1; Table S2). All collections were performed from late May to the end of October 2021. In addition, wild and cultivated cherry (*Prunus emarginata* [Douglas ex. Hook] [Rosaceae], *P. avium*) collections from three sites that followed methods for focal collection sites were included in the non-focal site analysis due to the emergence of several non-target *Drosophila* species other than *D. suzukii* (Table S2). Collected fruit from non-focal sites was incubated in collection containers and monitored according to the same procedure as outlined for focal sites at three laboratory sites: UBC (Vancouver, BC; temperature: 21 ± 2ºC), Summerland Research and Development Centre AAFC (Summerland, BC; 22 ± 2ºC), and the Ministry of Agriculture, Food, and Fisheries (Kelowna, BC; 20 ± 2ºC).

### Insect emergence and identification

Collected fruits were monitored three times per week for emergence of *Drosophila* species and parasitoids. Insects were mouth aspirated, collected into vials of 95% ethanol, and stored in labeled vials. *Drosophila* were sorted into *D. suzukii* and non-target *Drosophila* species, based on key characteristics described in Werner et al. (2020). Because (i) we collected fresh, undamaged fruit which is known to be infested almost exclusively by *D. suzukii*; (ii) fruit samples from which Drosophilidae species other than *D. suzukii* emerged were excluded from the analysis of focal sites; and (iii) the emerging parasitoids we recorded were all species previously known to parasitize *D. suzukii* (Abram et al. 2020; Abram et al. 2022), we made the simplifying assumption that any parasitoids emerging from the fruit samples likely emerged from *D. suzukii*. Although our method of rearing parasitoids and fly hosts from bulk fruit samples without separating out puparia is not ideal for determining conclusive host-parasitoid species associations (Abram et al. 2022), it is similar to the methodology used in previous studies of *D. suzukii* larval parasitism during foreign exploration in Asia (Daane et al. 2016; Girod et al. 2018a; Giorgini et al. 2019). Because of the large number of samples relative to the limited workforce involved in the study during a pandemic year, we also did not examine all material for unsuccessful parasitoid development or parasitoid diapause. However, careful inspection of unemerged pupae from a subset of samples (n=28; from 5 sites) from mid to late August revealed that only a small percentage (1.1%) of Drosophilidae pupae did not emerge; of these, 42% contained parasitoids that had partially or completely developed but had not emerged from the host puparium. Thus, our estimates of percent parasitism may be affected to a small degree by the fact that we did not include unsuccessful parasitoid development.

Parasitoids were identified using the identification key and notes provided in Abram et al. (2022). A representative sub-sample of Figitidae specimens were examined and identification confirmed by M. Buffington, while Braconidae were confirmed by R. Kula (USDA-ARS) and Pteromalidae by M. Gates (USDA-ARS). Voucher specimens are deposited in the National Insect Collection, National Museum of Natural History, Smithsonian Institution, Washington D.C.

### Data analysis

All counts of *Drosophila* and parasitoids were compiled and summary statistics were calculated for each host plant at the six focal sites. These included 1) total number of fruits, 2) total number of *D. suzukii* (including parasitized individuals) as estimated from the sum of emerged flies and parasitoids, 3) total percent parasitism estimated from the number of *D. suzukii* parasitized in all samples collected from a host plant relative to the total number of *D. suzukii* (including parasitized individuals), 4) minimum and maximum percent parasitism across sampling dates for each host plant site combination, and 5) percent parasitism separated by parasitoid species for each site and host plant. Our estimates of percent parasitism likely provide an underestimate of parasitism because eggs and small larvae present in fruit upon collection and incubation were not exposed to parasitism throughout their entire development.

To describe temporal characteristics of *D. suzukii* infestation and parasitism, we focused on *D. suzukii* and parasitoid emergence from the three plant species that we sampled at four or more focal sites, which also span the seasonal period during which *D. suzukii* reproduces on fruits in our study area: (1) Salmonberry, *R. spectabilis* (the earliest-season host infested by *D. suzukii* in our study area) which occurred at four focal sites; (2) Red elderberry, *S. racemosa* (a mid-season host for *D. suzukii*), which occurred at all six focal sites; and (3) Himalayan blackberry, *R. armeniacus* (the latest-season host for *D. suzukii*), which occurred at all six focal sites. We calculated the mean sampling duration (from first sampling date to final sampling date) for each host plant, the mean number of days until infestation levels (number of *D. suzukii* per berry) reached their maximum, days until maximum % parasitism, and number of days to first parasitism by *L. japonica* and *G. brasiliensis*. The latter three metrics were measured starting from the day *D. suzukii* first emerged, and were determined by pooling insect emergence numbers from the three pseudoreplicate samples by date for each host plant.

## Results

### Geographic occurrence

Both *L. japonica* and *G. brasiliensis* were found at all six focal sampling sites (total number of *D. suzukii* + parasitoids emerged = 13,923) and at 7 out of 14 non-focal sampling sites ((total number of *D. suzukii* + parasitoids emerged = 11,075) in the lower mainland/Fraser Valley region of British Columbia (Table 1; Table S1). Neither species of larval parasitoid was detected at non-focal sampling sites in the Okanagan Valley region (Figure 1), despite rearing a total of 1,278 *D. suzukii* from the fruit of nine different plant species, from 54 separate collecting events during the season (Table S2).

### Parasitoid-host plant associations

At the six focal sampling sites spanning a range of habitat types, both *L. japonica* and *G. brasiliensis* were found to be associated with *D. suzukii* throughout nearly the entire sampling period on many of the host plants we surveyed. Out of the total of 11 host plant species that were repeatedly sampled at these sites and from which *D. suzukii* emerged, *L. japonica* and *G. brasiliensis* were found to be associated with *D. suzukii* on 10 and 7 host plant species, respectively (Table 1).

Additional host plant associations for *L. japonica* and *G. brasiliensis* were observed in non-focal samples in the Lower Mainland region (Table S2). In these surveys, both *L. japonica* and *G. brasiliensis* were also found to be associated with *D. suzukii* on sour cherry (*P. emarginata*), sweet cherry (*P. avium*), and salal (*Gaultheria shallon* Pursh [Ericaceae]). *Ganaspis brasiliensis* was also reared from the fruits of two host plants in these samples from which it was not reared in focal surveys: highbush blueberry (*V. corymbosum*) and thimbleberry (*R. parvifolium*).

In addition to *L. japonica* and *G. brasiliensis*, we also found two other parasitoids uncommonly associated with *D. suzukii*. The larval parasitoid *Asobara* cf. *rufescens* (Förster) (Hymenoptera: Braconidae) was present at three focal sites in the Lower Mainland, associated with *D. suzukii* on raspberry (*R. idaeus*), Himalayan blackberry (*R. armeniacus*), and cultivated blackberry (*R. fruticosus*) (Table 1). The pupal parasitoid *Spalangia erythromera* Förster (Hymenoptera: Pteromalidae) was found associated with *D. suzukii* on Himalayan blackberry at one site on a single collection date (Table 1; Table S1).

### Seasonal occurrence of parasitoids, levels of parasitism, and parasitoid species composition

*Leptopilina japonica* and *G. brasiliensis* were associated with *D. suzukii* during most of the period when ripe fruit was sampled, from May to October (Table 1). *Drosophila suzukii* was first reared from fruit collected at a non-focal site on 23 May, *G. brasiliensis* on 28 May at a non-focal site, and *L. japonica* at a focal site on 5 June (Table S1). The last collection dates on which fruit yielded emerging *D. suzukii, G. brasiliensis*, and *L. japonica* were 16 October, 2 October, and 25 September, respectively (Table S1, Table S2).

Percent parasitism of *D. suzukii* at focal sampling sites was highly variable (Table 1). Total percent parasitism of *D. suzukii* by larval parasitoids across site-plant combinations ranged from 0-53%. Percent parasitism for individual site-plant-dates ranged from 0-66%. Notably, percent parasitism of *D. suzukii* was often highly variable over time within host plants at the same site: the mean difference between minimum and maximum percent parasitism within site-plants was 28.5 ± 3.7% (mean ± SE; n = 26). Among site-plants with parasitism, there was usually at least one sampling date with no parasitism at all (20/26 site-plants) (Table 1). Mean percent parasitism across all site-plants was 13.4 ± 2.3% (mean ± SE; n = 28 site-plants).

Of all of the parasitoids associated with *D. suzukii* at our focal sites (n = 2,755), roughly two thirds were *L. japonica* (67.2%) and one third *G. brasiliensis* (32.0%). Both *A*. cf. *rufescens* (0.6%) and *S. erythromera* (0.1%) were much less common (Table 1). While both *L. japonica* and *G. brasiliensis* were present at every focal site, parasitoid species composition among host plants and sampling dates was highly variable. *Leptopilina japonica* was the most commonly found parasitoid in 61.5% (16/26) of the host plant-site combinations with parasitism (Table 1). Both species were present in 61.5% (16/26) of host plant-site combinations with larval parasitism; *L. japonica* was the only larval parasitoid in 27.0% and *G. brasiliensis* was alone in the remaining 11.5%. Among host plants that were sampled at multiple focal sites, *L. japonica* appeared to be particularly dominant on salmonberry *R. spectabilis* (responsible for 83.4 ± 13.4% of all parasitism; n = 4 sites), and to a lesser extent on Himalayan blackberry *R. armeniacus* (64.1 ± 14.2%; n = 6 sites). In contrast, *G. brasiliensis* was clearly the dominant species on red elderberry *S. racemosa* (84.2 ± 7.1%; n = 6 sites).

Parasitoid species composition at non-focal sites in the lower mainland/Fraser Valley region (total n = 1,515 parasitoids) was 60.5% *L. japonica*, 38.2% *Ganaspis brasiliensis*, and 1.3% *Pachycrepoideus vindemiae* (Rondani) (Hymenoptera: Pteromalidae).

### Temporal dynamics of *D. suzukii* infestation and parasitism

For the three host plant species that were replicated across at least three sampling sites, we observed a range of temporal dynamics of *D. suzukii* infestation and parasitism with repeated sampling (Figure 2). For all three host plants, *D. suzukii* infestation levels reached their maximum level, on average, 16-19 days after fruit were first found to be infested with *D. suzukii* (Table 2). *Leptopilina japonica* and *G. brasiliensis* were detected, on average between 8 and 22 days after *D. suzukii* infestation was first detected, and percent parasitism peaked 19-33 days after the first *D. suzukii* infestation (Figure 2; Table 2). The delay between *D. suzukii* infestation and the first instance of parasitism was the longest for salmonberry for both *G. brasiliensis* and *L. japonica*, and was shortest for red elderberry and Himalayan blackberry for *G. brasiliensis* and *L. japonica*, respectively (Table 2). The delay until peak parasitism after the first instance of *D. suzukii* infestation was the longest for Himalayan blackberry and shortest for red elderberry (Table 2). There were no obvious temporal trends in parasitoid species composition (Figure 2), although there were trends in their relative dominance of each species on each plant overall (see above).

**Table 2.** Summary statistics describing temporal characteristics of *Drosophila suzukii* (*Ds*) infestation and parasitism by *L. japonica* (*Lj*) and *G. brasiliensis* (*Gb*) on three host plants that were sampled repeatedly (at least three times) at at least three sites: Salmonberry (*Rubus spectabilis*); Red elderberry, (*Sambucus racemosa*); and Himalayan blackberry (*Rubus armeniacus*). Refer to Figure 2 for a graphical representation of the temporal data.

| Time until event (days since first observed *Ds* infestation ± SE) | | | | | | |
| --- | --- | --- | --- | --- | --- | --- |
| Host plant | Number of sites | Average sampling period (days ± SE) | Max. *Ds* infestation | Max. % parasitism | First parasitism by *Lj* | First parasitism by *Gb* |
| Salmonberry<br>*Rubus spectabilis* | 4 | 27.7 ± 0.9 | 16.5 ± 4.8 | 22.75 ± 3.6 | 19.5 ± 3.3 | 22.0 ± 8.0 |
| Red elderberry<br>*Sambucus racemosa* | 6 | 25.8 ± 10.5 | 16.5 ± 6.7 | 19.0 ± 7.8 | 11.0 ± 4.5 | 10.5 ± 4.3 |
| Himalayan blackberry<br>*Rubus armeniacus* | 6 | 42.3 ± 17.3 | 19.0 ± 7.8 | 33.0 ± 13.5 | 8.4 ± 3.8 | 12.7 ± 5.2 |

**Figure 2.**
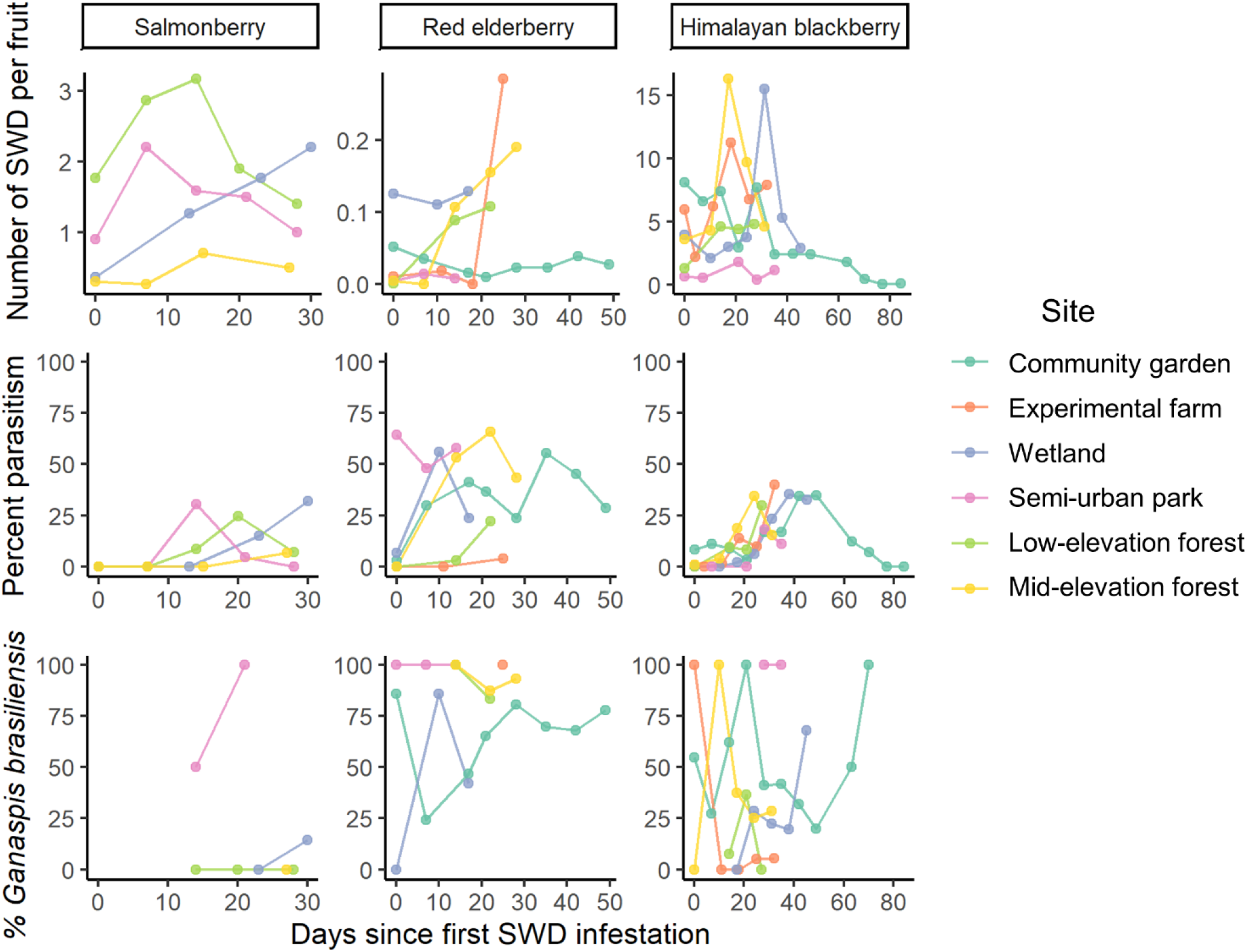
Temporal dynamics of the relative abundance of *D. suzukii* (SWD), total percent larval parasitism of SWD, and species composition of larval parasitoids (% *Ganaspis brasiliensis*) on three host plants that were sampled repeatedly (at least three times) at different sites: Salmonberry (*Rubus spectabilis*); Red elderberry,(*Sambucus racemosa*); and Himalayan blackberry (*Rubus armeniacus*). Time zero is set as the date at which SWD infestation was first observed on each host plant at each site.

## Discussion

Since first detecting the adventive presence of *L. japonica* and *G. brasiliensis* in south coastal British Columbia (Abram et al. 2020), the extent of their establishment and seasonal dynamics in association with *D. suzukii* has been unknown. In the present study, we clearly show that these two exotic parasitoids have reformed their close association with *D. suzukii* in this new geographic range and are now well established over the entire growing season across the landscape, in a relatively large geographic region in multiple habitats. We will now discuss the implications of our findings for the prospects of these parasitoids contributing to landscape-wide suppression of *D. suzukii* populations.

At our repeatedly surveyed sites in south coastal British Columbia, *Drosophila suzukii* was parasitized by one or both of *L. japonica* and *G. brasiliensis* at every site in most species of host plants that had *D. suzukii* infestation. While these sites represented a variety of habitat types with a considerable number of host plant species (e.g. semi-urban parks, wetlands, forests, experimental farms), they all shared the feature of being unmanaged habitats free of insecticide applications, or any other consistent disturbance events (e.g. harvesting of fruit). When *G. brasiliensis* and *L. japonica* were initially being considered for importation biological control of *D. suzukii*, it was anticipated that their main contribution to landscape-wide suppression of *D. suzukii* would be in these kinds of unmanaged habitats that could serve as sources or reservoirs for *D. suzukii* populations that disperse into commercial fruit crop fields on a seasonal or daily basis (Tonina et al. 2018; Urbaneja-Bernat et al. 2020). While our results show that these parasitoids are indeed parasitizing *D. suzukii* consistently in unmanaged habitats, they do not allow us to determine what their population-level impact will be on the landscape scale (see Abram et al. 2022). The net landscape-wide biological control effects of these parasitoids are yet to be determined and will depend on parasitism levels both in non-crop areas (this study), within crop fields (Abram et al., in prep.), and how these interact with the pest’s seasonal ecology, dispersal, and demography.

The extent to which the two species of larval parasitoids have recapitulated their ecology from their area of origin in their introduced range in North America is remarkable. They have evidently re-formed a close association with *D. suzukii* across a wide range of host plants including cultivated and wild shrubs, trees, and low-growing plants in a wide variety of habitats, and seem to be co-existing with each other in a manner very similar to their native range (Wang et al. 2020). Even the range of parasitism levels that we observed in our study (0-66%) and their level of variability, are qualitatively similar to those observed in Asia (Daane et al. 2016; Girod et al. 2018a; Giorgini et al. 2019). This implies that the parasitoids’ dispersal and host location strategies, and the ecological mechanisms allowing their co-existence, are at least to some degree generalizable to novel environments with different landscape features and plant communities. In addition, the two introduced parasitoids were responsible for nearly all of the parasitism we observed while native parasitoids of *Drosophila* are still, 11 years after the detection of *D. suzukii* in our study region, almost absent from the community parasitizing *D. suzukii* (with the possible minor parasitism by *A*. cf. *rufescens*, which may be of Palearctic origin, and *S. erythromera*, which is probably Holarctic). This finding highlights the fact that in the current global context of rapidly increasing biological invasions, unintentional introductions of exotic natural enemies (see Weber et al. 2021) that are well-adapted to exploiting their host can precede adaptation by native natural enemy communities.

An important finding of our study is that parasitism of *D. suzukii* by *L. japonica* and *G. brasiliensis* can be highly time-structured: in the early fruiting period of some host plants, parasitoids were often absent during the early stages of *D. suzukii* infestation, and the first incidence of parasitism was often not observed until ∼1-3 weeks after fruits were first infested with *D. suzukii*. One implication of this finding is that repeated sampling is necessary in order to detect the presence of these parasitoids, and accurately measure total parasitism levels of *D. suzukii* on a given host plant. For example, ‘point samples’ that happen to fall early in fruit ripening may yield false negatives with regards to parasitoid presence; this could partially explain their apparent absence at some of our non-focal sites in the Lower Mainland (Figure 1). Similarly, point samples late in the infestation cycle may overestimate the number of *D. suzukii* killed by parasitism. A second implication of this finding is that the *D. suzukii* that are the first to disperse to colonize a new host plant’s ripening fruit may escape parasitism, creating a series of partial spatiotemporal ‘refuges from parasitism’ over the landscape and season that could potentially diminish the long-term population-level biological control impacts of parasitoids (Hawkins et al. 1993). To date, however, the majority of theoretical work on refuges has focused on how they might promote the stability of host-parasitoid dynamics, rather than their impact on pest suppression (Mills and Getz 1996; Takagi 1999). Highly time-structured host or prey mortality resulting from spatiotemporal refuges created by host and natural enemy dispersal among patches has long been a topic of biological control and population ecology research (e.g. Huffaker 1958; reviewed in Takagi 1999) and may be critical to consider when forecasting prospects for the success of biological control of *D. suzukii* by larval parasitoids.

The pest status of *D. suzukii* in North America has been at least partially attributed to release from specialized natural enemies present in its area of origin that it escaped when it invaded North America and Europe (e.g., Asplen et al. 2015; Gabarra et al. 2015). Now that it has been reunited with its two most closely associated larval parasitoids, will its populations be suppressed enough to impact pest management? This is currently a difficult question to answer, as the pest status of *D. suzukii* in Asia where *L. japonica* and *G. brasiliensis* originate appears to be regionally variable (Lee et al. 2011) and the role of these larval parasitoids in suppressing its populations there is still not well understood. What is clear is that *D. suzukii* is currently still highly abundant in our study region (Table 1), despite the apparent ubiquity of the two parasitoids in unmanaged habitats and the highest parasitism rates of *D. suzukii* yet measured outside of their area of origin. However, the current study is just a ‘snapshot’ of a single year, less than five years since the first detection of the two parasitoid species in the study area (*L. japonica* - 2016; *G. brasiliensis* -2019). Thus, parasitism levels measured here may still be in a state of flux, as stable host population suppression by introduced natural enemies can often take several years (e.g., Caltagirone 1981). In addition, future studies will need to make use of updated methods (Abram et al. 2022) for measuring larval parasitism of *D. suzukii* and determining its relationship to population-level impact on the pest on a landscape scale.

## Acknowledgements

We thank Jade Sherwood, Mairi Robertson, and Tyler Nelson for assistance with field collections and Nathan Earley for his comments on the manuscript. We thank Markus Clodius and Seth Nussbaum for access to unsprayed berry fields. We also thank Robert Kula (USDA-ARS) for providing expert identification of Braconidae parasitoids and Mike Gates (USDA-ARS) for identification of Pteromalidae. Mention of trade names or commercial products in this publication is solely for the purpose of providing specific information and does not imply recommendation or endorsement by the U.S. Department of Agriculture. USDA is an equal opportunity employer and provider.

## Funding

T.H. and S.A. were supported in part by the Canadian Agriculture Partnership, a federal-provincial-territorial initiative, and the Lower Mainland Horticultural Improvement Association. This research (funding to P.K.A, C.E.M., and J.C.) is part of Organic Science Cluster 3, led by the Organic Federation of Canada in collaboration with the Organic Agriculture Centre of Canada at Dalhousie University, supported by Agriculture and Agri-Food Canada’s Canadian Agricultural Partnership - AgriScience Program. P.K.A. and C.E.M. were also supported by funding from Agriculture and Agri-Food Canada, A-BASE #2955.

## Notes

### Competing Interest Statement

The authors have declared no competing interest.

### Summary of Updates

Small typos and textual edits have been made since the original posting.

